# Exchange of metabolic pathway intermediates as the origin for cross-feeding interactions in *E. coli* auxotrophic co-cultures

**DOI:** 10.64898/2026.10.08.757639

**Authors:** Yijing Cai, Maciek R. Antoniewicz

## Abstract

A large fraction of microbes in natural environments are auxotrophs that rely on metabolite exchange for growth. The Black Queen hypothesis and related frameworks explain gene loss and reliance on shared metabolites. Yet it is not fully understood how metabolite cross-feeding emerges and gives rise to experimentally observable community performance. Here, we propose that exchange of biosynthetic pathway intermediates provides a mechanistic route for the emergence of metabolite cross-feeding. Building on prior work demonstrating intermediate exchange in *E. coli* auxotrophs, we investigated co-cultures between a methionine auxotroph (ΔmetA) and a panel of histidine auxotrophs. Among these, only ΔhisD exhibited efficient co-culture growth with ΔmetA, suggesting a unique role for the final step of histidine biosynthesis, which converts histidinol to histidine. We found that ΔmetA-ΔhisD co-cultures accumulated high levels of extracellular histidinol (>200 µM), suggesting a mechanism for metabolite exchange. Spent media from these co-cultures supported growth of all histidine auxotrophs except ΔhisD, confirming that histidinol, rather than histidine, was the dominant exchanged metabolite. Furthermore, we observed that ΔmetA efficiently converted histidinol to histidine only under methionine-limited conditions, suggesting that nutrient limitation promoted overflow of converted histidine to the extracellular environment. To quantitatively assess whether histidinol exchange could account for the observed co-culture behavior, we constructed a mechanistic model that links extracellular histidinol dynamics to co-culture growth through independently measured conversion capacity of the ΔmetA strain. Rather than fitting growth empirically, we used experimentally derived parameters to predict co-culture growth from the measured histidinol-to-histidine conversion flux. The model accurately recapitulated the timing and magnitude of growth and population dynamics, supporting the idea that accumulation of histidinol enabled the conversion-driven supply of histidine that sustained ΔhisD, while coupling growth between the two strains. Taken together, these results suggest that metabolite cross-feeding in syntrophic co-cultures can emerge from the accumulation of pathway intermediates and their conditional conversion under nutrient limitation, providing a plausible mechanism for the rapid establishment of mutualistic interactions.

## 1. Introduction

In microbial communities, acquiring nutrients from the surrounding environment is essential for survival. A large fraction of naturally occurring microbes cannot synthesize one or more essential metabolites and instead rely on neighbors to supply them^1–3^. This dependence is not rare: recent genomic analyses indicate that more than 70% of bacteria in natural communities are auxotrophic for at least one amino acid^1,4^. The prevalence of auxotrophy has been attributed to the selective advantage of shedding costly biosynthetic functions, a principle formalized in the Black Queen Hypothesis, which proposes that cells gain fitness by exploiting public goods that leak from other community members^5,6^. Experimental evolution has shown that prototrophic *E. coli* can evolve complete amino acid dependency in fewer than 2,000 generations^7^, confirming that metabolic dependencies arise readily once cross-feeding opportunities exist.

The cross-feeding interactions that sustain these auxotrophic communities are increasingly recognized as a key force shaping community structure, stability, and function^8–10^. Yet a central question remains open: how and why does the cross-feeding of amino acids, costly biosynthetic building blocks, originate? The release of metabolic waste products such as acetate and hydrogen is well understood as a consequence of overflow metabolism and growth-rate optimization^11,12^. Similarly, the release of incompletely metabolized resources can be explained by the growth-rate advantage of partial metabolism^13^. However, as Chacón et al.^14^ and Harcombe et al.^15^ have noted, neither of these frameworks explains why bacteria release amino acids, the very molecules they need to grow. The mechanisms proposed for amino acid externalization, passive diffusion, cell lysis, phage-mediated release, and active export, remain debated^16,17^, and the question of how amino acid cross-feeding becomes established in the first place has been identified as a key gap in the field^15,18^.

*E. coli* auxotroph co-cultures have become the primary experimental model for studying cross-feeding interactions. In 2010, Wintermute and Silver screened 46 auxotrophic *E. coli* mutants from the Keio single-gene knockout collection^19^ in all 1,035 pairwise combinations and found that 17% showed positive co-growth, establishing a widespread capacity for metabolite exchange^20^. Mee and colleagues extended this to communities of up to 14 amino acid auxotrophs, revealing numerous syntrophic interactions with both positive and negative epistasis^21^. These studies, together with the demonstration that cross-feeding pairs can outcompete wild-type prototrophs^22^, have motivated a substantial body of work on the ecology and evolution of amino acid cross-feeding.

Recent work from our laboratory revealed that the exchange of biosynthetic pathway intermediates, rather than end-product amino acids, is a prevalent and previously underappreciated feature of auxotrophic co-cultures. Hong et al.^23^ showed that in co-cultures of auxotrophic *E. coli* sharing the same amino acid requirement, strains that should not be able to help each other if only amino acids are exchanged, 23 out of 68 pairs showed positive growth, indicating that pathway intermediates were being exchanged. In the histidine pathway, histidinol was identified and verified as the cross-fed intermediate. The possibility that pathway intermediates could serve as exchange currencies has also been raised in other contexts. Mee and colleagues noted that when two strains’ biosynthetic routes intersect at a shared intermediate, the strains more readily divide the pathway for mutual benefit^21^. In endosymbiotic systems, pathway splitting at intermediate hand-off points is well documented, the tryptophan biosynthetic pathway in the *Cinara cedri* aphid consortium is partitioned between *Buchnera* and *Serratia*, with a pathway intermediate as the exchanged species^24,25^, and kinetic modeling of such consortia has shown that the splitting point corresponds to the most membrane-permeable intermediate^26^. Despite these observations, while the occurrence of intermediate exchange has been noted, no study has examined how it functionally contributes to cross-feeding performance, nor has any co-culture model incorporated the bidirectional feedback loop through which intermediate accumulation and condition-dependent conversion reinforce each other. The computational frameworks most widely used to model co-culture dynamics including COMETS^27^, OptCom^28^, SteadyCom^29^, and dynamic FBA approaches^30^, define exchangeable metabolites primarily as end products and treat exchange kinetics as fixed parameters, independent of partner physiology. As a result, neither the role of pathway intermediates as exchange currencies nor the self-reinforcing feedback between mutual nutrient limitation and intermediate conversion is represented in current models.

Here, we address the gaps using the ΔhisD–ΔmetA co-culture as a model system. We demonstrate that intermediate exchange confers a distinct growth advantage to ΔhisD over other histidine auxotrophs in co-culture and establish the mechanism underlying this advantage: accumulation of the pathway intermediate histidinol in the shared environment, followed by its conversion to histidine by ΔmetA through a process that is gated by nutrient limitation. Together, these elements form a self-reinforcing exchange loop that we capture in a mechanistic model parameterized from independent measurements. The model reproduces co-culture dynamics quantitatively, providing evidence that intermediate exchange driven by this feedback loop is sufficient to explain the system’s behavior. More broadly, our findings offer a new explanation for how cross-feeding can emerge: gene-loss-driven intermediate accumulation followed by the co-culture environment activates conversion by the partner, and the resulting feedback loop establishes a stable mutualistic interaction.

## 2. Results

### 2.1. ΔhisD shows a growth advantage among histidine auxotrophs in co-culture

To determine whether disrupting different steps of the histidine biosynthetic pathway differentially affects co-culture performance, we co-cultured the methionine auxotroph ΔmetA^31^ with a series of histidine auxotrophs (ΔhisI, ΔhisA, ΔhisF, ΔhisB, ΔhisC, and ΔhisD) in minimal medium. In each pair, the histidine auxotroph can synthesize methionine but not histidine, while ΔmetA can synthesize histidine but not methionine, setting up a potential reciprocal exchange.

The histidine biosynthetic pathway and the position of each lesion are shown in Figure 1A^32,33^. In the final step, hisD encoded histidinol dehydrogenase catalysis the NAD-dependent oxidation of histidinol to histidine is disturbed in ΔhisD. It was previously shown to be histidinol is the only histidine intermediate that can be cross-fed between ΔhisDand other histidine auxotrophs^23^.

**Figure 1.**
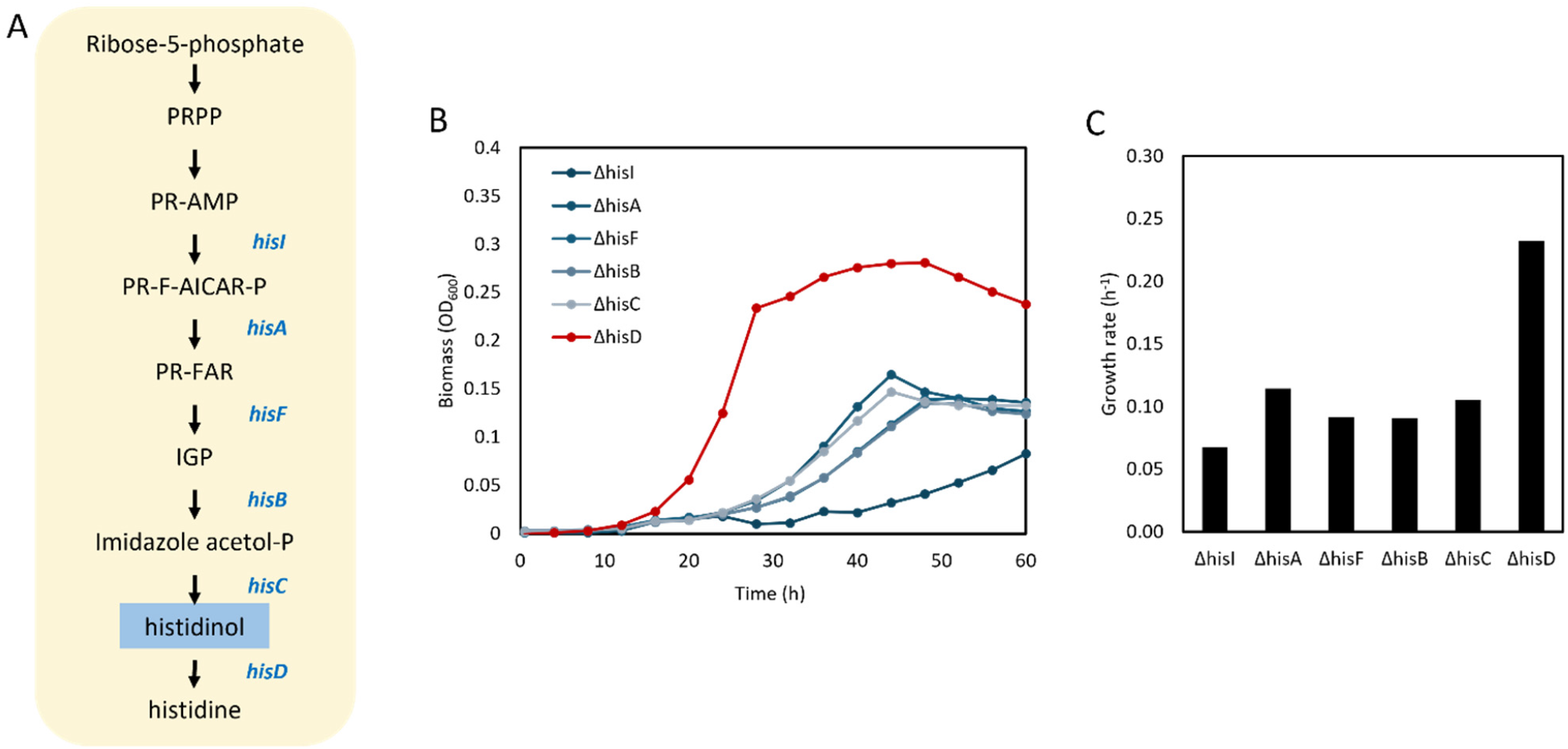
ΔhisD shows a unique growth advantage among histidine auxotrophs in co-culture. (A) The histidine biosynthetic pathway with gene names and the position of each knockout. Histidinol (highlighted) is the first dephosphorylated intermediate. (B) Growth curves of ΔmetA co-cultured with each histidine auxotroph in M9 minimal medium over 60 hours. ΔhisD (red) reaches approximately 0.28 OD_600_, roughly two-fold higher than all other pairs. (C) Growth rate (h^−1^) for each co-culture pair. ΔhisD–ΔmetA shows a growth rate of approximately 0.23 h^−1^, more than double the other histidine auxotroph pairs.

Growth of each co-culture was monitored over 60 hours (Figure 1B). Most histidine knockouts behaved similarly, reaching a final biomass of approximately 0.10–0.13 OD_600_ with growth rates near 0.07–0.10 h^−1^ (Figure 1C). The ΔhisD–ΔmetA co-culture grew markedly faster, reaching a peak OD_600_ of approximately 0.28 and a growth rate of approximately 0.23 h^−1^ (Figure 1C), roughly two-fold higher than the other pairs. This phenotype was reproducible across biological replicates and was not observed in monocultures, indicating that the enhanced growth arises from strain interaction rather than from intrinsic differences.

These results suggest that disruption at the position in histidine biosynthesis where histidinol is the exchangeable between cells plays an important role in the performance of cross-feeding between the two auxotrophs.

### 2.2 Histidinol accumulates extracellularly in the co-culture

Because ΔhisD is blocked at the oxidation of histidinol to histidine, flux through the upstream pathway continues could leads to histidinol accumulates. We therefore hypothesized that histidinol, rather than histidine, accumulates in the ΔhisD–ΔmetA co-culture and mediates cross-feeding.

Direct GC-MS profiling of histidinol proved unreliable under our analytical conditions. Histidinol carries three polar, reactive functional groups, including an imidazole nitrogen, which complicates complete derivatization and reliable detection. Histidine itself is recognized as problematic by GC-MS for the same structural reasons, and no validated direct chromatographic or mass-spectrometric assay for histidinol has been established^34–37^. A 1 mM histidinol standard did not yield a consistent, identifiable peak. We therefore developed a functional bioassay, using the growth of a histidine auxotroph as a quantitative reporter of histidinol, an approach validated previously for amino acid biosensing^38^.

Supernatant from ΔhisD–ΔmetA co-cultures at late exponential growth phase was harvested by centrifugation, filtered to remove cells, supplemented with glucose, and supplied as growth medium to individual histidine auxotrophs (Figure 2A). If histidinol accumulates, spent medium should support the growth of histidine auxotrophs other than strain ΔhisD due to absence histidinol dehydrogenase. The result was consistent with this prediction. Spent medium supported robust growth of ΔhisI, ΔhisA, ΔhisF, ΔhisB, and ΔhisC, whereas ΔhisD showed negligible growth (Figure 2B). Because ΔhisD cannot use histidinol but can use free histidine, its failure to grow on spent medium confirms that the accumulated species is histidinol rather than histidine, and that growth is not supported by some unrelated nutrient. Parallel cultures on defined medium supplemented with 100 µM histidinol reproduced the same pattern (Figure 2B, blue bars), confirming that the bioassay reports histidinol specifically.

**Figure 2.**
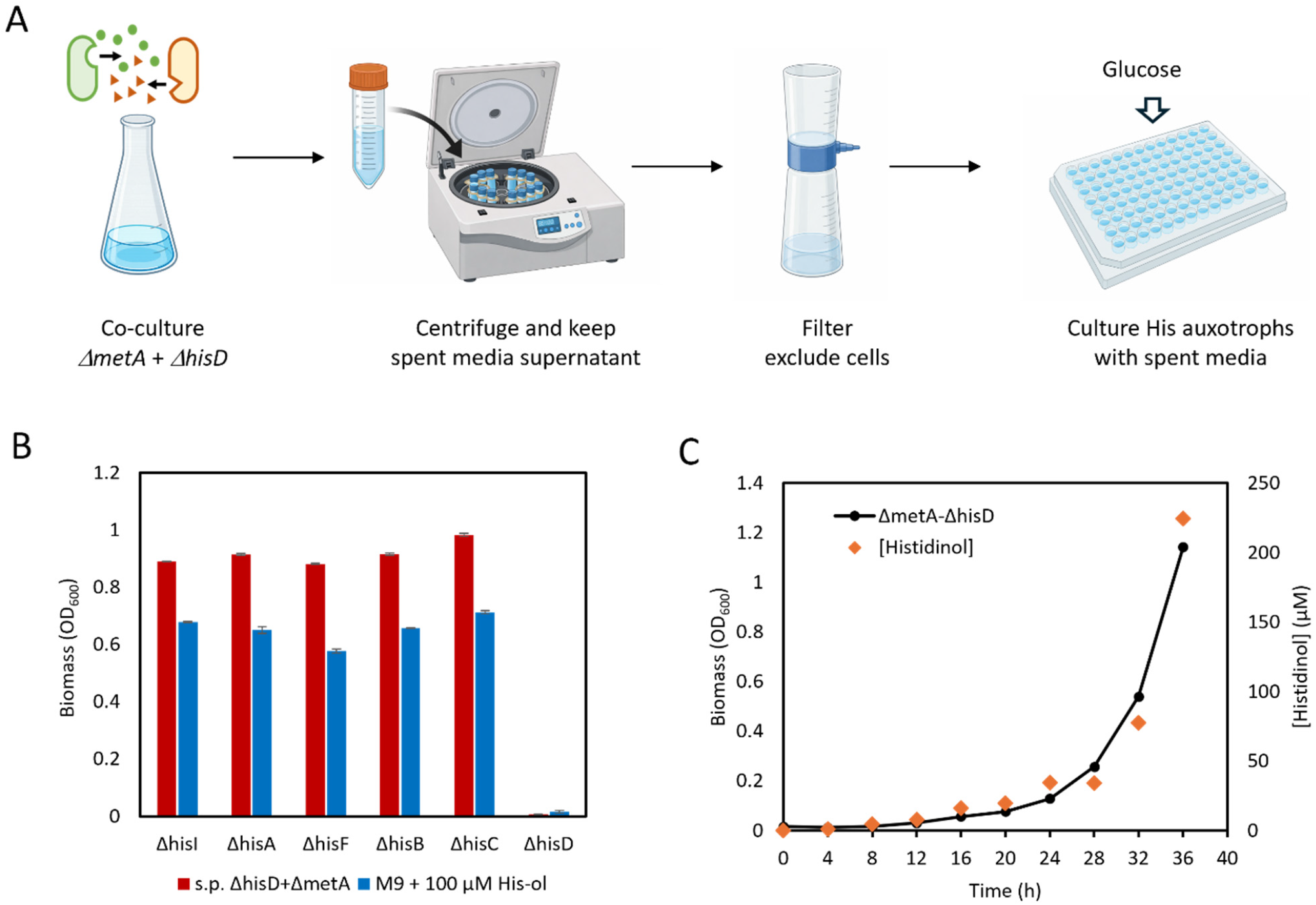
Histidinol accumulates to >200 µM in the ΔhisD–ΔmetA co-culture. (A) Spent media bioassay workflow: ΔhisD–ΔmetA co-culture supernatant is harvested by centrifugation, filtered to exclude cells, supplemented with glucose, and used to culture individual histidine auxotrophs. (B) Bioassay results. Spent medium from ΔhisD–ΔmetA co-cultures (red bars) supports growth of all histidine auxotrophs except ΔhisD, confirming that the accumulated species is histidinol, not histidine. Defined medium with 100 µM histidinol (blue bars) reproduces the same pattern. (C) Time course of co-culture growth (black circles, left axis) and extracellular histidinol concentration (orange diamonds, right axis) quantified by growth-based bioassay. Histidinol reaches approximately 220 µM at late exponential phase.

To quantify histidinol, we established a calibration curve using ΔhisA grown in minimal medium across a histidinol gradient. Final biomass was linear with histidinol concentration between 0 and 40 µM (OD_600_ = 0.0094 × [His-ol], R^2^ = 0.9962; Supplementary Information S1), so biomass serves as a quantitative proxy within this range. Applying this calibration to spent-medium samples showed that extracellular histidinol accumulated progressively during co-culture, reaching approximately 220 µM at late exponential phase (Figure 2C).

Together, these results demonstrate that histidinol accumulates to physiologically significant concentrations and is the primary species sustaining the growth of histidine auxotrophs in this system.

### 2.3 Nutrient limitation enables conversion of histidinol to histidine in ΔmetA

Histidinol accumulation alone does not explain the enhanced co-culture growth, because the partner that consumes histidinol, ΔmetA, must convert it to histidine efficiently enough to feed ΔhisD in return. We therefore measured histidine production by ΔmetA across a histidinol gradient under two nutrient conditions.

ΔmetA was grown in minimal medium with 1 mM methionine (nutrient-replete condition) and supernatant was harvested at OD_600_ ≈ 0.7. Histidine was quantified with a calibration curve from ΔhisD grown across a histidine gradient, which excludes any contribution from residual histidinol; ΔhisD biomass was linear with histidine concentration (OD_600_ = 0.0152 × [His], R^2^ = 0.9961; Supplementary Information S1). Under methionine-replete conditions, ΔmetA produced essentially no histidine when supplied with 1–100 µM histidinol, and only about 3.5 µM histidine even at 500 µM histidinol (Figure 3, left panel). At co-culture-relevant histidinol concentrations, this conversion is far too low to support the growth observed.

**Figure 3.**
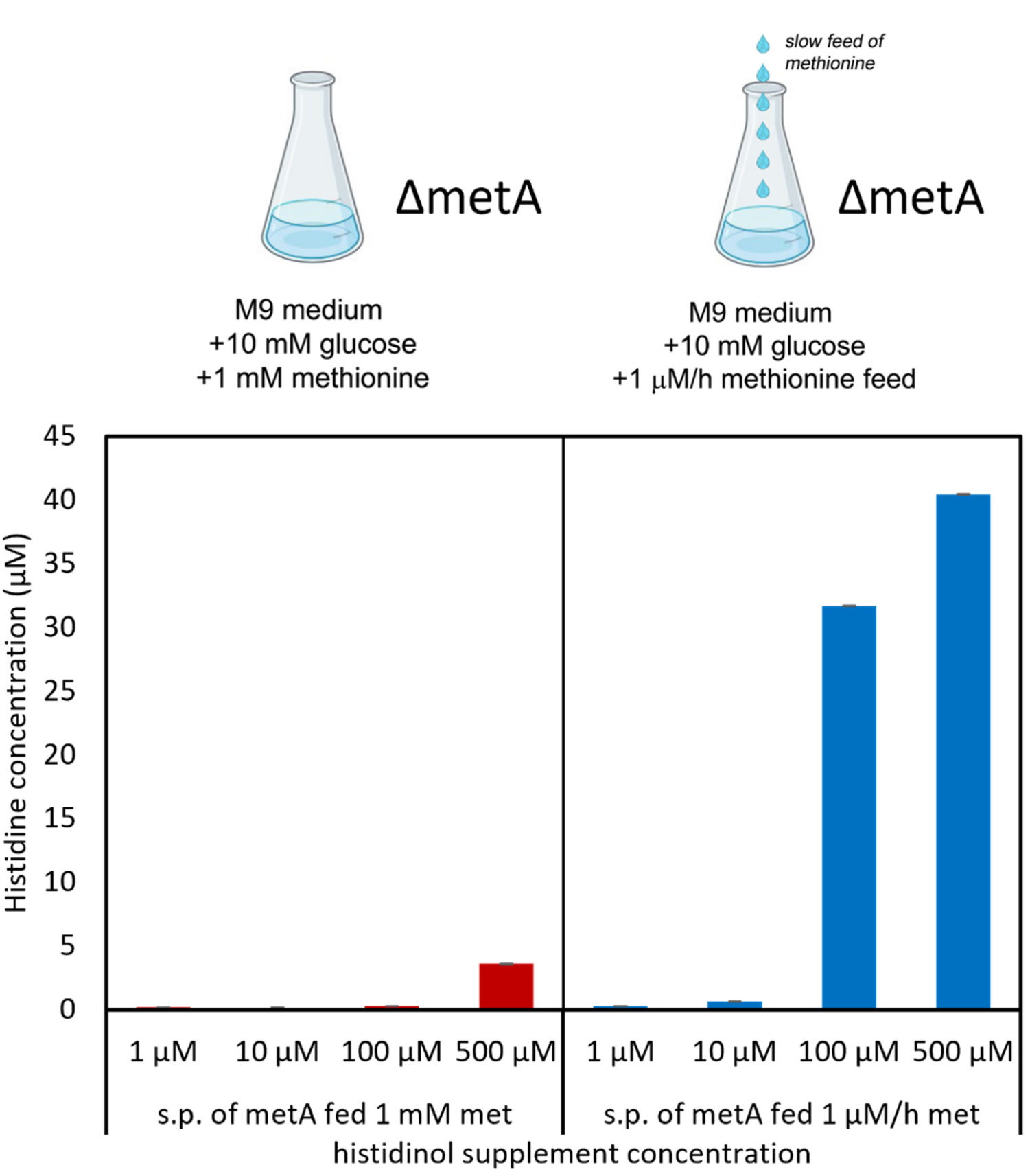
Nutrient limitation activates histidinol-to-histidine conversion by ΔmetA. ΔmetA was cultured under methionine-replete (1 mM methionine, left) or methionine-limited (1 µM h^−1^ feed, right) conditions. Supernatant was harvested and supplemented with histidinol at concentrations from 1–500 µM. Histidine produced was quantified using a ΔhisD growth-based bioassay. Under replete conditions, ΔmetA converts essentially no histidinol (red bars). Under methionine limitation, conversion increases dramatically (blue bars), reaching approximately 32 µM histidine from 100 µM histidinol.

To test whether nutrient limitation changes histidinol conversion, we repeated the experiment while supplying methionine at a constant rate of 1 µM/h, holding ΔmetA in a steady methionine-limited state that mimics the co-culture environment. Conversion increased dramatically. With 100 µM histidinol, methionine-limited ΔmetA produced approximately 32 µM histidine, and produced approximately 40 µM at 500 µM histidinol (Figure 3, right panel). This level of histidine production is sufficient to support ΔhisD growth to the levels observed in co-culture.

These results identify nutrient limitation as a switch that converts ΔmetA from a poor to an effective histidinol consumer. Critically, this switch is self-activating: in co-culture, ΔmetA is necessarily methionine-limited because ΔhisD supplies methionine only at a rate sufficient for slow growth. The co-culture environment itself enables productive intermediate conversion and create a feed-back loop of histidinol to histidine from strain ΔhisD to ΔmetA and back to ΔhisD.

### 2.4 A dynamic exchange model reproduces growth and composition dynamics

To test whether histidinol exchange and reciprocal methionine support are together sufficient to explain the co-culture behavior, we built a dynamic model with three state variables: ΔmetA biomass (*X*_*M*_), ΔhisD biomass (*X*_*D*_), and extracellular histidinol (*Hol*). The model structure is illustrated schematically in Figure 4A. The initial composition was set to the measured value at inoculation, and biomass fractions were allowed to evolve.

**Figure 4.**
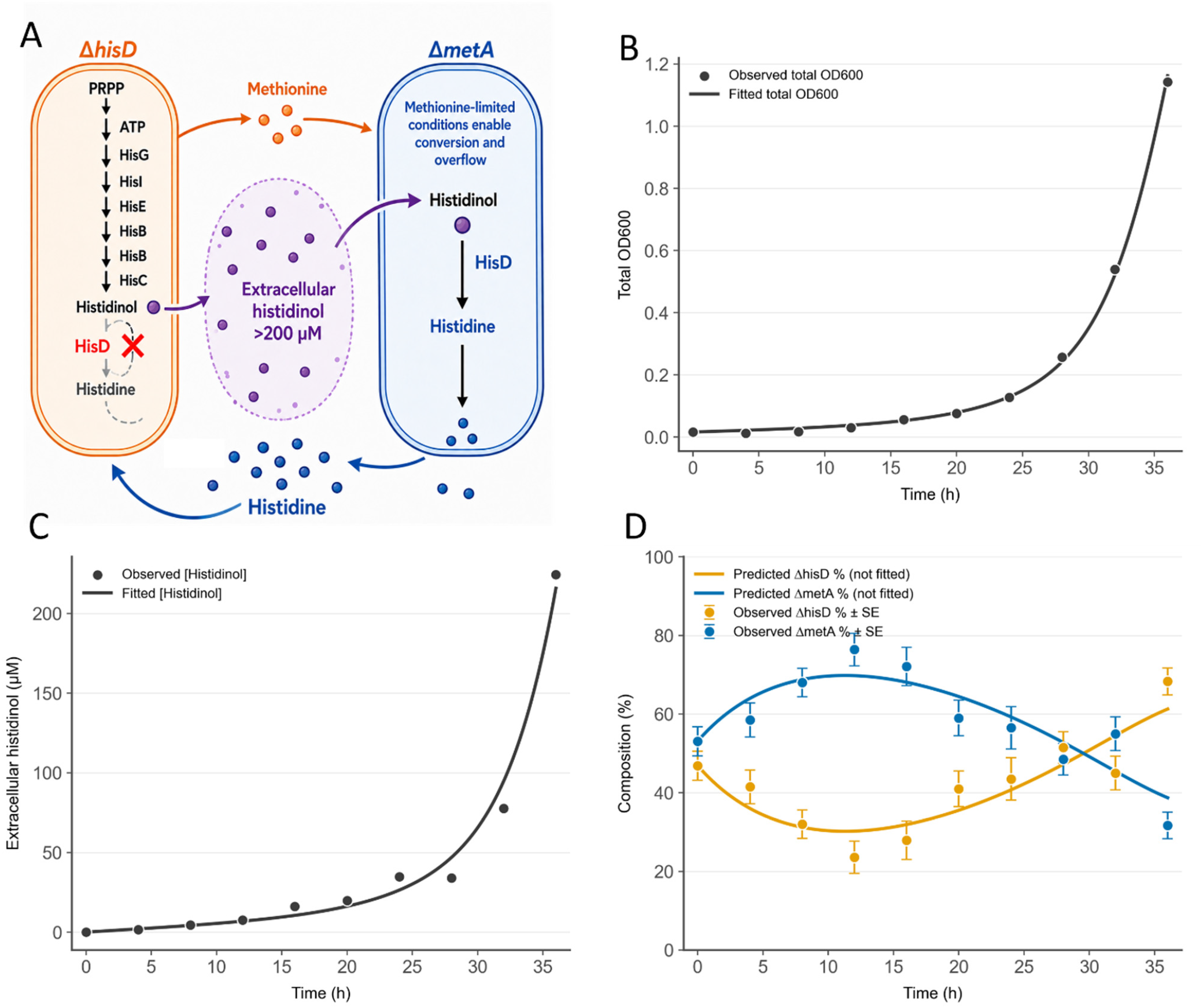
A dynamic exchange model constrained by independent measurements reproduces co-culture dynamics. (A) Schematic of the exchange model. ΔhisD produces histidinol, which accumulates extracellularly. ΔmetA converts histidinol to histidine under methionine-limited conditions and feeds histidine back to ΔhisD. Methionine flows from ΔhisD to ΔmetA. (B) Total biomass: model prediction (line) vs. observed data (dots); R^2^ = 0.999. (C) Extracellular histidinol concentration: R^2^ = 0.984. (D) Strain composition: predicted (lines) vs. observed (dots with error bars); R^2^ = 0.797. Composition was not included in the fitting objective.

Histidinol-to-histidine conversion by ΔmetA was described by a saturating rate law:

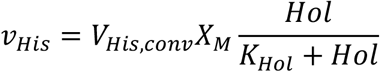

Because ΔmetA is methionine-limited throughout co-culture, *V*_*His,conv*_, represents the maximum conversion rate from histidinol to histidine catalyzed by per biomass ΔmetA, was fixed at the experimentally measured value of 10.45 µM h^−1^ OD^−1^ of ΔmetA under nutrient-limited conditions. *K*_*Hol*_ represents the half-saturation constant. Methionine support from ΔhisD to ΔmetA, *v*_*Met*_, was described as:

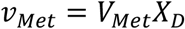

*V*_*Met*_ represents the production rate of methionine per biomass ΔhisD. Histidinol production by

ΔhisD and the balance of extracellular histidinol *v*_*Hol*_ were described by:

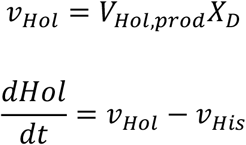

*V*_*Hol,prod*_ represents the production rate of histidinol per biomass ΔhisD. Biomass growth of each strain was linked to the exchanged metabolite it receives, subject to a carrying-capacity term:

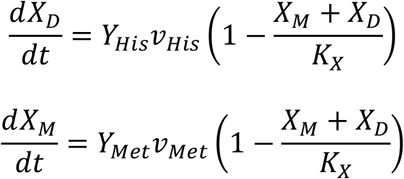

Three parameters were measured independently and fixed during fitting: *V*_*His,conv*_ = 10.45 µM h^−1^ OD^−1^ from the histidinol-conversion experiment under limitation (Supplementary Information S3); *Y*_*His*_ = 0.13 OD µM^−1^ from histidine-supplementation growth curves; and *Y*_*Met*_ = 0.19 OD µM^−1^ from methionine-supplementation growth curves (Supplementary Information S2). The remaining parameters were optimized against the time courses of total biomass and extracellular histidinol only. Strain composition was not included in the fitting objective and instead served as an independent validation of the model.

The model reproduced both the growth trajectory (R^2^ = 0.999) and the accumulation of extracellular histidinol (R^2^ = 0.985) (Figure 4B, C). Most informatively, the model also reproduced the non-monotonic strain composition (R^2^ = 0.803; Figure 4D), despite composition not being included in the fitting objective. ΔmetA dominated early, when little histidinol had yet accumulated, and ΔhisD overtook it after roughly 20 hours, once histidinol accumulated to levels where conversion by methionine-limited ΔmetA became substantial and histidine feedback to ΔhisD accelerated its growth.

The model is constrained by independently measured conversion parameters, and fitted only to total biomass and histidinol concentration, recovers the composition reversal as an emergent prediction strongly supports the claim that histidinol exchange gated by ΔmetA’s nutrient limitation, together with reciprocal methionine support, is sufficient to generate the observed co-culture dynamics.

### 2.5 Fitted parameters are physically interpretable

The four fitted parameters are all physically interpretable and consistent with independent observations. The optimized parameters were *V*_*Hol,prod*_ = 66.87 µM h^−1^ OD^−1^, *V*_*Met*_ = 0.92 µM h^−1^ OD^−1^, *K*_*Hol*_ = 255.48 µM, and *K*_*X*_ = 2.39 OD.

The histidinol production rate, *V*_*Hol,prod*_ = 66.87 µM h^−1^ OD^−1^, is approximately 40-fold larger than the methionine supply rate, *V*_*Met*_ = 0.92 µM h^−1^ OD^−1^. This asymmetry is expected: ΔhisD produces histidinol as a direct consequence of its gene knockout, which blocks the terminal oxidation while upstream flux continues unabated, resulting in massive intracellular accumulation and overflow. In contrast, methionine release by ΔhisD is not driven by a pathway block; it reflects the small, constitutive leakage that all cells exhibit. The very low *V*_*Met*_ also explains why we were unable to detect extracellular methionine by either GC-MS or bioassay, despite repeated attempts.

The half-saturation constant for histidinol conversion, *K*_*Hol*_ = 255.48 µM, is substantially higher than the published Michaelis constant of histidinol dehydrogenase (HisD), which ranges from 11–18 µM in kinetic measurements of the purified enzyme^32,39^. This discrepancy is understandable, unlike the enzyme assay, whole-cell conversion requires histidinol/histidine to cross the membrane twice to be absorbed and secreted from ΔmetA. So the apparent half-saturation constant reflects the combined resistance of transport and catalysis rather than enzyme kinetics alone.

The carrying capacity, *K*_*X*_ = 2.39 OD, is consistent with the maximal biomass we routinely observe for *E. coli* in M9 minimal medium with glucose as the sole carbon source. *K*_*X*_ represents the total population density at which growth ceases due to resource depletion (carbon, nitrogen, or other essential nutrients), and its fitted value falls within the range expected for this medium.

That each fitted parameter independently aligns with known biochemistry indicates the model captures the mechanistic architecture of the co-culture rather than merely fitting its trajectory, lending confidence that the intermediate exchange feedback loop it describes reflects the actual basis of co-culture performance.

## 3. Discussion

In this study, we investigated how intermediate exchange drives co-culture performance in auxotrophic *E. coli* co-cultures. As the model system we used the ΔhisD–ΔmetA co-culture, where ΔhisD is auxotrophic for histidine and ΔmetA is auxotrophic for methionine. We demonstrated that ΔhisD exhibits a roughly two-fold growth advantage over other histidine auxotrophs in co-culture with ΔmetA, that the pathway intermediate histidinol accumulates extracellularly to over 200 µM, and that ΔmetA converts histidinol to histidine efficiently only under the methionine-limited conditions that co-culture itself imposes. A dynamic exchange model parameterized from independently measured values reproduced the dynamic of strain composition. Taken together, these results demonstrate that intermediate exchange, gated by mutual nutrient limitation, provides a sufficient mechanism for the emergence of amino acid cross-feeding from a single gene loss event.

An important finding in this study was that the availability of histidinol alone does not predict productive cross-feeding. When ΔmetA was grown in the presence of excess methionine, it converted even abundant histidinol to histidine poorly, and the reciprocal feedback to ΔhisD failed. Only when ΔmetA was methionine-limited, the state it necessarily occupies in co-culture, did conversion rise to a level sufficient to sustain its partner. These results have an important implication: cross-feeding is a conditional process gated by partner physiology rather than a static property of the metabolite. This feedback loop is what closes the exchange and stabilizes the interaction. Interestingly, this is consistent with our recent ^13^C-MFA study of the ΔmetA strain^40^, which revealed that methionine limitation induces widespread metabolic rewiring, including increased TCA cycle flux, activation of dormant pathways, and secretion of specific amino acids. The enhanced histidinol-to-histidine conversion we observe here likely reflects a similar nutrient-limitation-induced metabolic state. Yet the mechanism underlying this enhanced conversion remains unknown. Elucidating this mechanism represents a valuable direction for future work, as it would clarify whether condition-dependent conversion is a general feature of nutrient-limited auxotrophs or specific to this system, and could inform the rational design of synthetic consortia that exploit limitation-activated exchange.

The roughly two-fold growth advantage of ΔhisD over the upstream histidine knockouts is best understood as a division of labor within a single biosynthetic pathway rather than exchange of a finished product. ΔhisD completes most of the reactions of histidine biosynthesis and hands ΔmetA a nearly finished substrate; ΔmetA contributes only the final step before returning histidine. The community splits one pathway across two genomes, with histidinol as the hand-off point. This is far more economical than the alternative available to the upstream knockouts, in which ΔmetA must run the entire histidine pathway de novo to supply a partner that contributes nothing to histidine synthesis. The shared-intermediate principle proposed by Mee and colleagues^21^ is realized here in concrete form. In metabolic engineering, pathway splitting across co-culture strains has been employed primarily to reduce metabolic burden and relieve end-product feedback inhibition^41–43^, and the exchange of intermediates between strains is typically viewed as an unavoidable cost, a source of dilution and carbon loss that must be minimized^43^. Our results suggest a complementary design principle: intermediate accumulation driven by a pathway truncation can itself serve as the productive mechanism for cross-feeding, and the nutrient status of the consuming partner can act as a natural valve that controls conversion efficiency. Engineering co-cultures that deliberately exploit gene-knockout-driven intermediate overflow and condition-dependent conversion, rather than treating these as liabilities, may open new strategies for constructing stable and self-regulating synthetic consortia.

The dynamic exchange model we constructed provided additional insights. The model’s principal value is that a small number of physiologically interpretable parameters, with the key conversion capacity measured independently, suffice to explain the emergent behavior. That the model reproduces the composition reversal, ΔmetA dominance early, ΔhisD overtaking later, without fitting to composition data is a particularly strong validation, because composition dynamics reflect the interplay of all exchange fluxes and growth rates simultaneously. The four fitted parameters are all consistent with independent observations: *V*_*Hol,prod*_ is approximately 40-fold larger than *V*_*Met*_, matching the expectation that gene-knockout-driven overflow vastly exceeds constitutive leakage; *K*_*Hol*_is substantially higher than the published *K*_*M*_ of purified histidinol dehydrogenase^32,39^, consistent with the additional transport resistance of whole-cell conversion; and *K*_*X*_ matches the typical carrying capacity of *E. coli* in M9 glucose medium. That each parameter independently aligns with known biochemistry indicates that the model captures the mechanistic architecture of the co-culture rather than merely fitting its trajectory. Still, the model also has clear limits. The methionine exchange rate was inferred from fitting rather than measured directly, because extracellular methionine remained below detection limits. Integrating this tractable kinetic framework with genome-scale community models that capture whole-cell metabolic rerouting, connecting, for example, the condition-dependent conversion we observe to the broader flux rewiring identified in our recent ^13^C-MFA study of methionine-limited ΔmetA ^40^ would be a valuable direction for future work.

Insights from this study also have implications for understanding how obligate metabolic dependencies arise in nature. Under the Black Queen Hypothesis, selection favors loss of costly biosynthetic functions whose products are available as public goods^5^, and auxotrophs can gain a direct fitness advantage when the missing metabolite is supplied^6^. Our results suggest that gene loss at a biosynthetic step resulting in the accumulation of intermediate is a particularly attractive evolutionary entry point. It is genomically cheap, a single gene inactivation, yet it relieves the cost of the blocked reaction, and because the truncated pathway leaks a usable intermediate, the loss simultaneously creates the public good a partner can exploit. The self-reinforcing nutrient-limitation feedback we describe could then stabilize the nascent interaction, since the partners are driven into the cooperative regime exactly as they begin to limit one another. This offers a plausible route from a chance gene loss to a stable obligate mutualism without requiring the prior evolution of export machinery, consistent with the observed stability of obligate cross-feeding after gene loss^22^ and with models in which interdependencies arise from individually advantageous reductions in metabolic burden^44^. These findings also connect to our recent observation that ΔmetA secretes glutamate, threonine, and glycine under methionine limitation^40^, raising the possibility that nutrient-limitation-induced metabolite release is a general feature that facilitates the establishment of cross-feeding interactions across multiple metabolic pathways simultaneously.

Overall, this study demonstrates that the emergence of amino acid cross-feeding does not require the evolution of dedicated secretion, transport, or cooperative regulation. Instead, it can arise from three elements that are inherent consequences of gene loss in a shared environment: intermediate accumulation, condition-dependent conversion, and a self-reinforcing feedback loop driven by mutual nutrient limitation. Clearly, the conventional view of auxotrophic cross-feeding as a simple exchange of finished amino acids is incomplete. Our results show that in order to understand the true dynamics of auxotrophic co-cultures, it is important to consider the identity of the actually exchanged metabolite, the physiological state of the partner, and the feedback between the two. To advance understanding of microbial communities, it will be important to test whether this intermediate exchange mechanism generalizes across other biosynthetic pathways, to resolve the molecular basis of limitation-activated conversion, and to incorporate condition-dependent intermediate exchange into community metabolic models.

## 4. Materials and methods

### 4.1. Strains and materials

All auxotrophic *E. coli* strains were obtained from the Keio knockout collection (GE Healthcare Dharmacon), which were generated by one-step gene inactivation in *E. coli* K-12 BW25113 (Baba et al., 2006). Chemicals and M9 minimal medium were purchased from Sigma-Aldrich (St. Louis, MO).

### 4.2. Co-culture of auxotrophic strains

Precultures were prepared in M9 medium supplemented with 1 mM glucose, 3% (v/v) LB medium, and 50 µg/mL kanamycin, and incubated overnight at 37 °C with shaking. For co-culture experiments, 100 µL of each preculture (ΔmetA paired with ΔhisI, ΔhisA, ΔhisF, ΔhisB, ΔhisC, or ΔhisD) was inoculated into 10 mL of M9 minimal medium containing 10 mM glucose and 50 µg/mL kanamycin in 150 mL baffled Erlenmeyer flasks at 37 °C with shaking. Growth was monitored by measuring optical density at 600 nm (OD_600_) using a spectrophotometer (Eppendorf BioPhotometer). Growth rates were calculated from the slope of ln(OD_600_) versus time during the exponential growth phase. For the ΔhisD–ΔmetA co-culture, supernatant samples were collected and filtered (0.22 µm) at each time point for quantification of histidinol by bioassay. At late exponential phase, the entire culture volume was harvested by centrifugation and filtration through a 0.22 µm filter for spent media characterization.

### 4.3. Histidinol to histidine conversion characterization

To characterize the conversion of histidinol to histidine by ΔmetA under different nutrient conditions, ΔmetA precultures were inoculated into 10 mL of M9 minimal medium containing 10 mM glucose and 50 µg/mL kanamycin. For the nutrient-replete condition, medium was supplemented with 1 mM methionine. For the nutrient-limited condition, methionine was supplied hourly at 100 µL of a 100 µM methionine solution, corresponding to a feeding rate of 1 µM h^−1^. In both conditions, histidinol was supplemented at concentrations of 1, 10, 100, or 500 µM. At late exponential phase, spent media were harvested and filtered (0.22 µm). Histidine concentration in the spent media was quantified using the ΔhisD-based bioassay described below. Samples were stored at −20 °C if not analyzed immediately.

### 4.3. Bioassay calibration curves

For histidinol quantification, ΔhisA was cultured in M9 minimal medium containing 10 mM glucose, 50 µg/mL kanamycin, and histidinol at concentrations spanning 0–40 µM for 24 h. Final biomass was plotted against histidinol concentration, yielding a linear relationship (OD_600_ = 0.0094 × [His-ol], R^2^ = 0.9962; Supplementary Information S1). For histidine quantification, ΔhisD was used in the same procedure with a histidine concentration gradient (OD_600_ = 0.0152 × [His], R^2^ = 0.9961).

### 4.3. Spent media characterization

To determine the identity of the accumulated species in co-culture spent media, precultures of individual histidine auxotrophs (ΔhisI, ΔhisA, ΔhisF, ΔhisB, ΔhisC, and ΔhisD) were inoculated into co-culture spent media supplemented with 10 mM glucose in 96-well plates and cultured for 24 h at 37 °C, with OD_600_ recorded every 30 min. For quantification of histidinol or histidine in spent media, ΔhisA or ΔhisD was cultured in spent media supplemented with 10 mM glucose for 24 h at 37 °C. Concentrations were calculated by matching final biomass to the corresponding calibration curves described above.

### 4.4. Direct measurements of model parameters

Biomass yield coefficients for histidine (*Y*_*His*_) and methionine (*Y*_*Met*_) were determined from endpoint growth experiments. ΔhisD was cultured in M9 minimal medium containing 10 mM glucose, 50 µg/mL kanamycin, and histidine at 0, 10, 20, 50, 100, or 200 µM. ΔmetA was cultured under the same conditions with methionine at 0, 10, 20, 50, or 100 µM. Cultures were grown in 96-well plates for 24 h at 37 °C and endpoint OD_600_ was recorded. Plate reader absorbance values were converted to spectrophotometer-equivalent OD_600_ using a linear calibration determined from parallel measurements. Final biomass was fitted to a saturating yield model.

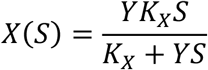

*Y* is the yield coefficient and *K*_*X*_ is the carrying capacity. The fitted values were *Y*_*His*_ = 0.13 OD µM^−1^ and *Y*_*Met*_ = 0.19 OD µM^−1^ (Supplementary Information S2).

To determine the apparent histidinol-to-histidine conversion rate (*V*_*His,conv*_), ΔmetA was cultured in M9 minimal medium containing 10 mM glucose, 1 mM methionine, and 50 µg/mL kanamycin until OD_600_ reached approximately 0.7. Cells were harvested by centrifugation, washed three times with M9 medium to remove residual methionine, and resuspended in M9 medium containing 10 mM glucose and 200 µM histidinol. Supernatant samples were collected and filtered at 0, 2, 5, and 12 hours, and histidine concentration was quantified using the ΔhisD bioassay described in Section 4.6. The apparent conversion rate was determined from a linear fit of histidine concentration versus time (slope = 7.85 µM h^−1^, R^2^ = 0.989), normalized to the measured biomass of the ΔmetA culture (OD_600_ = 0.751), yielding *V*_*His,conv*_ = 10.5 µM h^−1^ OD^−1^ (Supplementary Information S3).

## Author Statement

Yijing Cai: Conceptualization, Validation, Formal analysis, Investigation, Data Curation, Roles/Writing - original draft. Maciek R. Antoniewicz: Conceptualization, Data Curation, Writing - Review & Editing, Supervision, Project administration, Funding acquisition.

