## Supplemental Information for "Exchange of metabolic pathway intermediates as the origin for cross-feeding interactions in *E. coli* auxotrophic co-cultures"

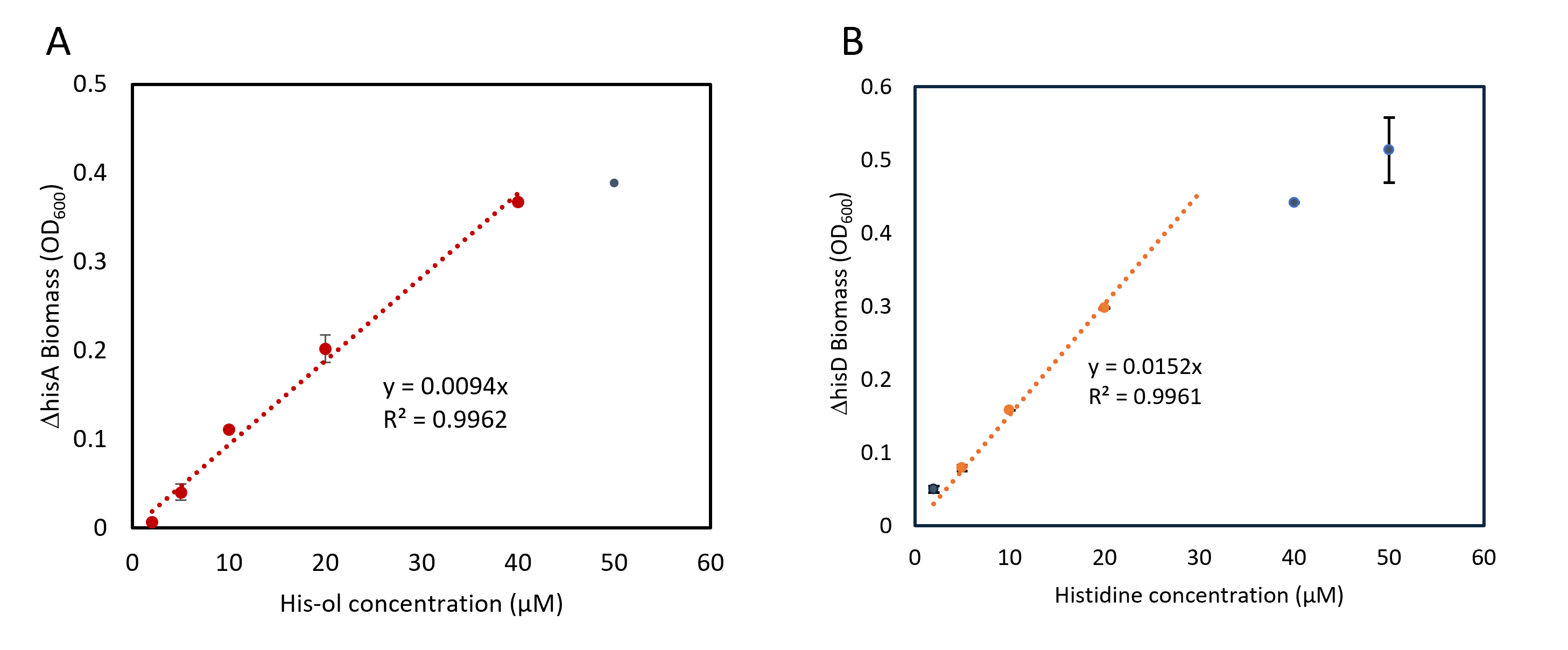
Supplementary Information

**Figure S1. Bioassay calibration curves for histidinol and histidine quantification.** (A) ΔhisA grown in M9 minimal medium supplemented with histidinol at concentrations spanning 0–50 µM for 24 h. Endpoint biomass (OD₆₀₀) was linear with histidinol concentration up to 40 µM (y = 0.0094x, R² = 0.9962; red, linear range used for calibration). The data point at 50 µM (gray) deviates from linearity and was excluded from the calibration. (B) ΔhisD grown under the same conditions with histidine at concentrations spanning 0–50 µM. Endpoint biomass was linear with histidine concentration up to 40 µM (y = 0.0152x, R² = 0.9961; orange, linear range). Data points at 40 and 50 µM (gray) begin to saturate. Error bars represent standard deviation (n = 2).

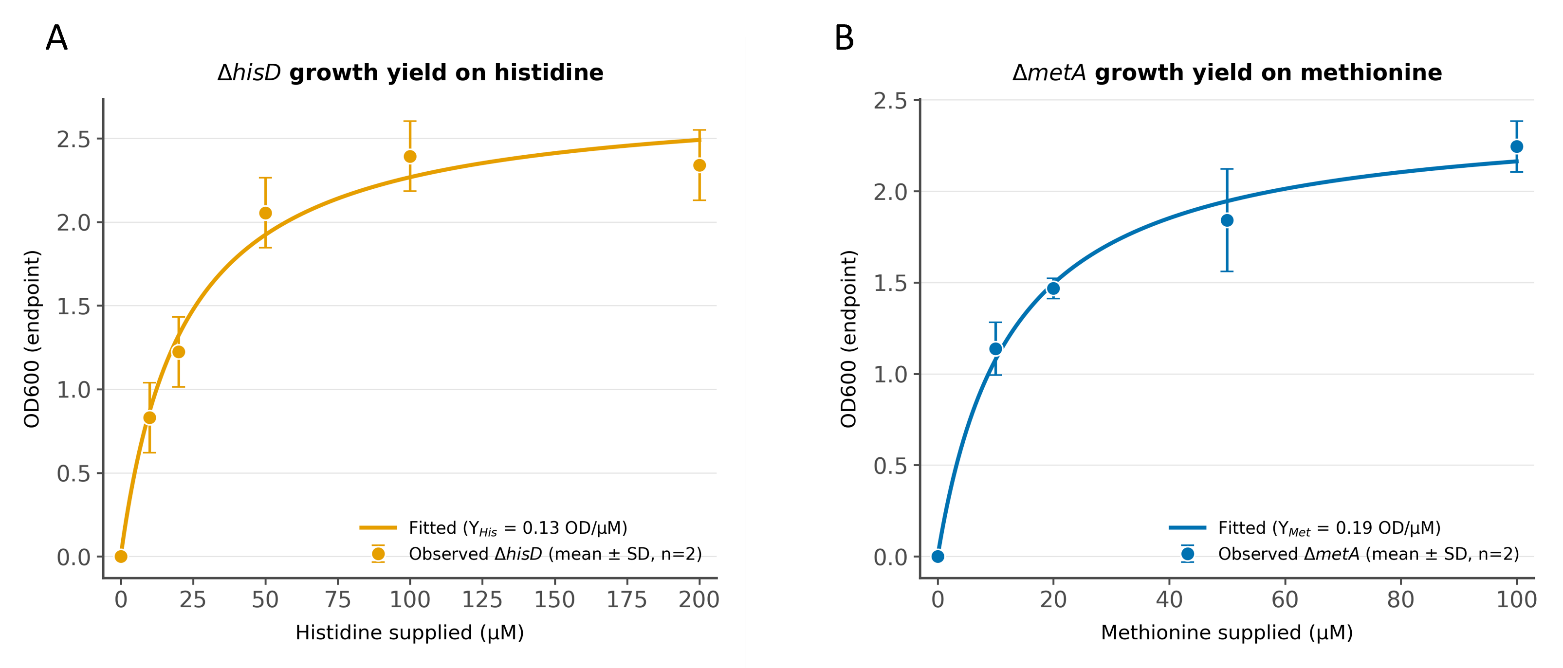

**Figure S2. Growth yield determination for** $Y_{His}$ and $Y_{Met}$. (A) ΔhisD endpoint biomass (OD₆₀₀) as a function of histidine supplied (0–200 µM). Data were fitted to a saturating yield model X(S) = Y · K_X · S / (K_X + Y · S), yielding $Y_{His}$ = 0.13 OD µM⁻¹. (B) ΔmetA endpoint biomass as a function of methionine supplied (0–100 µM), yielding $Y_{Met}$= 0.19 OD µM⁻¹. Plate reader absorbance values were converted to spectrophotometer-equivalent OD₆₀₀ using a linear calibration. Error bars represent standard deviation (n = 2). The yield coefficient Y represents the initial slope of the dose-response curve (the low-concentration linear regime), and $K_{X}$represents the carrying capacity at saturation.

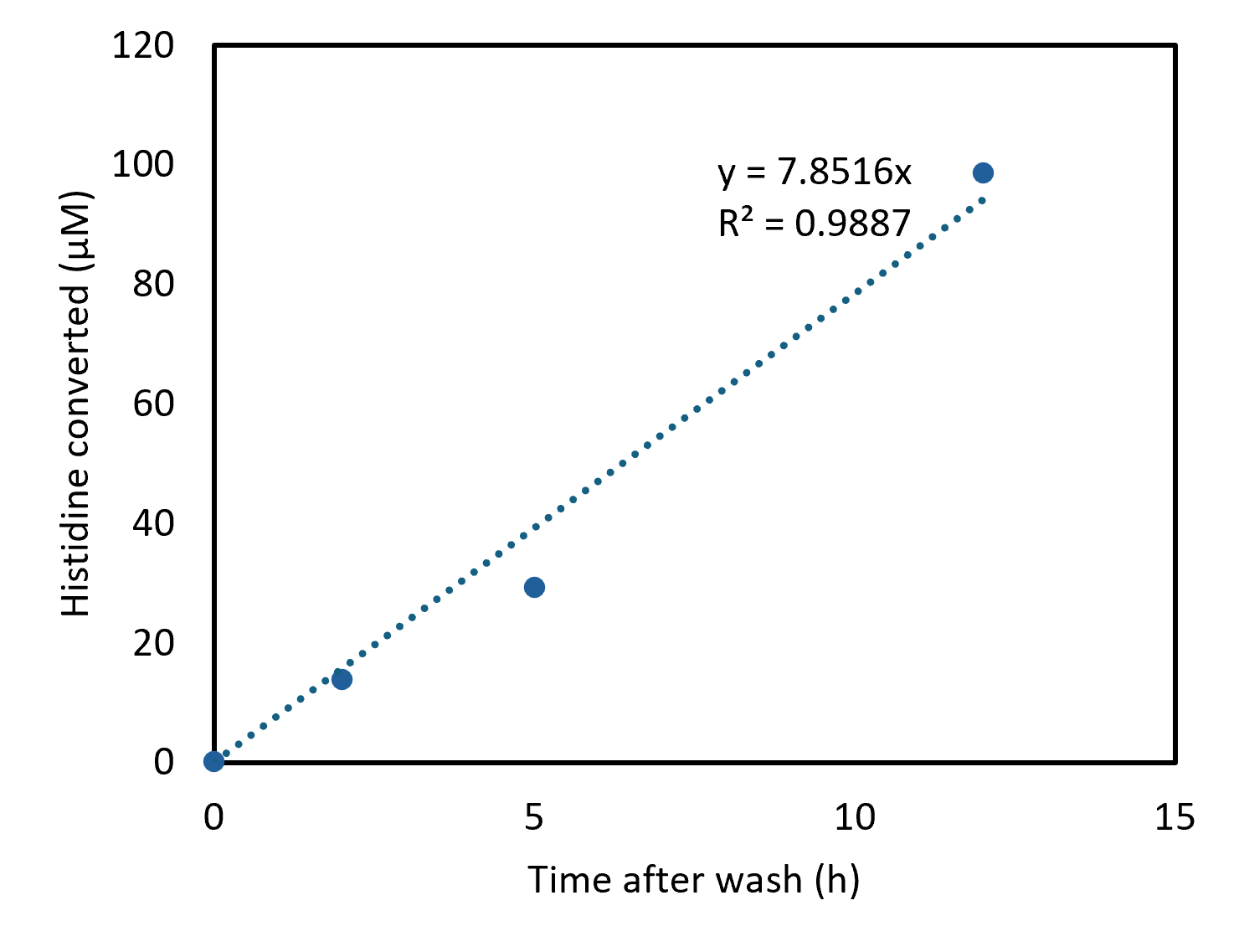

**Figure S3. Direct measurement of the apparent histidinol-to-histidine conversion rate (**$V_{His, conv}$**).** ΔmetA was cultured with 1 mM methionine to OD₆₀₀ ≈ 0.75, washed three times with M9 medium, and resuspended in M9 medium containing 10 mM glucose and 200 µM histidinol. Supernatant samples were collected at 0, 2, 5, and 12 h and histidine concentration was quantified using the ΔhisD bioassay (Figure S1B). Histidine accumulated linearly over time (y = 7.85x, R² = 0.989). Normalizing to the measured biomass (OD₆₀₀ = 0.751) yields $V_{His, conv}$= 10.5 µM h⁻¹ OD⁻¹, which was used as a fixed parameter in the dynamic exchange model.
